# Alignment of virus-host protein-protein interaction networks by integer linear programming: SARS-CoV-2

**DOI:** 10.1101/2020.07.07.191247

**Authors:** Mercè Llabrés, Gabriel Valiente

## Abstract

Beside socio-economic issues, coronavirus pandemic COVID-19, the infectious disease caused by the newly discovered coronavirus SARS-CoV-2, has caused a deep impact in the scientific community, that has considerably increased its effort to discover the infection strategies of the new virus. Among the extensive and crucial research that has been carried out in the last few months, the analysis of the virus-host relationship plays an important role in drug discovery. Virus-host protein-protein interactions are the active agents in virus replication, and the analysis of virus-host protein-protein interaction networks is fundamental to the study of the virus-host relationship. We have adapted and implemented a recent integer linear programming model for protein-protein interaction network alignment to virus-host networks, and obtained a consensus alignment of the SARS-CoV-1 and SARS-CoV-2 virus-host protein-protein interaction networks. Despite the lack of shared human proteins in these virus-host networks and the low number of preserved virus-host interactions, the consensus alignment revealed aligned human proteins that share a function related to viral infection, as well as human proteins of high functional similarity that interact with SARS-CoV-1 and SARS-CoV-2 proteins, whose alignment would preserve these virus-host interactions.

## 1 Introduction

The present outbreak of a coronavirus-associated acute respiratory disease, the COVID-19 pandemic, has forced the scientific community to rapidly analyse the virus-host relationships of the new coronavirus (SARS-CoV-2) human infection. Thus, in less than a month, several databases as [1–3] have been created to collect all SARS-CoV-2 and COVID-19 information, and the SARS-CoV-2-human protein-protein interaction network was built [4]. As stated in [5], “The *Coronaviridae Study Group* (CSG) of the International Committee on Taxonomy of Viruses […] has assessed the placement of the human pathogen, tentatively named 2019-nCoV, within the *Coronaviridae*. Based on phylogeny, taxonomy and established practice, the CSG recognizes this virus as forming a sister clade to the prototype human and bat severe acute respiratory syndrome coronaviruses (SARS-CoVs) of the species *Severe acute respiratory syndrome-related coronavirus*, and designates it as SARS-CoV-2.” Therefore, the closest known human pathogen to SARS-CoV-2 is the coronavirus SARS-CoV that appeared in 2003 [6], also called SARS-CoV-1.

Understanding the mechanism of the SARS-CoV-2 infection is a crucial step towards the discovery of antiviral drugs and vaccines. The *modus operandi* of every viral infection is through the interaction between viral proteins and host proteins, in order to use the host cells to replicate. In this line of research, virus-host protein-protein interaction networks, a particular form of protein-protein interaction networks, have become appropriate to analyse virus-host relationships, and information on well-known and studied virus-host protein-protein interaction networks can be carried over to new ones by way of protein-protein interaction network comparison and alignment. See [7, 8] for comprehensive reviews.

The general problem of protein-protein interaction network alignment has been explored in the last two decades, and several tools have been already proposed and implemented [9–14]. However, the particular case of virus-host protein-protein interaction network alignment problem has not been fully studied yet.

We have recently developed a compact reformulation of a quadratic programming model for the protein-protein interaction network alignment problem as an integer linear program, which has been proven to be suitable for the alignment of virus-host protein-protein interaction networks [15]. Our proposed model can be solved using state-of-the-art mathematical modeling software such as AMPL [16] and integer linear programming software tools such as IBM ILOG CPLEX Optimization Studio and Gurobi Optimizer. In this work, we adapt and implement a modification of the aforementioned alignment method to align the virus-host protein-protein interaction networks of SARS-CoV-1 and SARS-CoV-2, in order to elucidate information on the infection mechanism of SARS-CoV-2 based on current knowledge on the infection mechanism of SARS-CoV-1.

## 2 Materials and methods

In the integer linear programming formulation of the protein-protein interaction network alignment problem, a virus-host protein-protein interaction network is represented by an undirected bipartite graph *G* = (*U, V, E*), with a node *u* ∈ *U* for each virus protein, a node *v* ∈ *V* for each host protein, and an edge {*u, v*} ∈ *E* for each virus-host protein-protein interaction. Let *G* = (*U, V, E*) and *G*′ = (*U*′, *V* ′, *E*′) be the two virus-host protein-protein interaction networks to be aligned, and let *A* = (*a*_*ij*_) and *B* = (*b*_*k*𝓁_) be their weighted adjacency matrices, where the weight of an entry *a*_*ij*_ ∈ [0, 1] is the confidence score of the interaction {*i, j*} ∈ *E*, and the weight of an entry *b*_*k*𝓁_ ∈ [0, 1] is the confidence score of the interaction {*k, f*} ∈ *E*′. Let also *S* = (*s*_*ik*_) be a similarity matrix between the nodes of the two networks, with each *s*_*ik*_ ∈ [0, 1] the similarity score of *i* ∈ *U* ∪ *V* and *k* ∈ *U*′ ∪ *V* ′.

Let us define a binary variable *x*_*ik*_ for each *i* ∈ *U* ∪ *V* and each *k* ∈ *U*′ ∪ *V* ′, where *x*_*ik*_ = 1 if node *i* of the first network is aligned with node *k* of the second network, and *x*_*ik*_ = 0 otherwise. Then, an alignment of two virus-host protein-protein interaction networks *G* = (*U, V, E*) and *G*′ = (*U*′, *V* ′, *E*′) is represented by the binary matrix *X* = (*x*_*ik*_). Let us also define an integer variable *y*_*ik*_ for each *i* ∈ *U* ∪ *V* and each *k* ∈ *U*′ ∪ *V* ′, where each integer variable *y*_*ik*_ is intended to represent

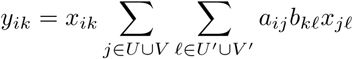

for *i* ∈ *U* ∪ *V* and *k* ∈ *U*′ ∪ *V* ′. In this way, if *x*_*ik*_ = 0, *y*_*ik*_ = 0, and if *x*_*ik*_ = 1, *y*_*ik*_ is the weight of those edges incident to node *i* in *G* that are preserved by the alignment.

Then, the goal of the integer linear programming model is to maximize

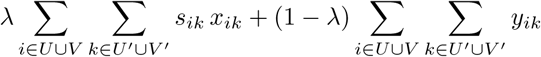

subject to the constraints

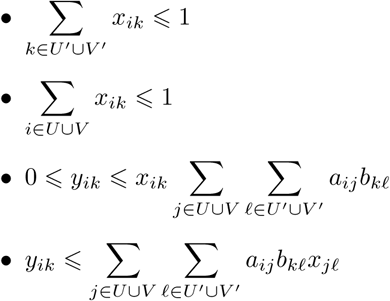

for *i* ∈ *U* ∪ *V* and *k* ∈ *U*′ ∪ *V* ′, where *λ* is a parameter, with 0 ⩽ *λ* ⩽ 1, to control the balance between protein similarity scores and protein-protein interaction weights: only node scores are considered when *λ* = 1, and only edge scores are taken into account when *λ* = 0.

It is easy to see that this integer linear programming formulation is equivalent to the integer quadratic programming formulation of the network alignment problem given in [9]. In fact, the previous constraints entail

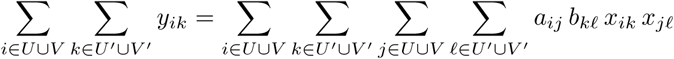

The objective function comes from the PathBLAST [11] idea that protein-protein network alignment be based on a log-probability-like criterion, with matching terms corresponding to both proteins and interactions [9]. The first sum in the objective function,

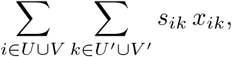

represents the global similarity of the aligned proteins, while the second sum,

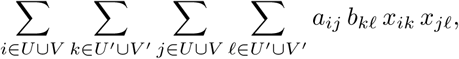

represents the weight of those edges that are preserved by the alignment; that is, those pairs of edges (*i, j*) ∈ *E* and (*k*, 𝓁) ∈ *E*′ such that node *i* is aligned with node *k* and node *j* is aligned with node 𝓁.

Now, since we are aligning virus-host protein-protein interaction networks, we add the constraints

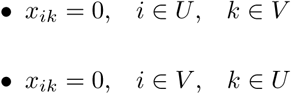

to ensure that viral proteins are aligned with viral proteins, and that host proteins are aligned with host proteins.

Let *m* = |*U*| + |*V* | and *n* = |*U*′| + |*V* ′|. The resulting integer linear programming formulation of the virus-host protein-protein network alignment problem has *O*(*mn*) binary variables, integer variables, and constraints.

## 3 Results and discussion

There are 130 interactions between 29 SARS-CoV-1 proteins and 109 human proteins in the March 2020 release of the VirHostNet database [1], as well as 332 interactions between 26 SARS-CoV-2 proteins and 332 human proteins from [4] in release 4.2.13 of the IntAct database [2]. Thus, the SARS-CoV-1-Human network has 138 nodes and 130 edges, while the SARS-CoV-2-Human network has 358 nodes and 332 edges. Notice that only 6 of these 109 and 332 human proteins (P27448, Q5JRX3, Q7KZI7, Q9BW92, Q9H4F8,and Q9Y6E2) interact with both SARS-CoV-1 and SARS-CoV-2 proteins.

We have obtained the amino acid sequences for the SARS-CoV-1 and human proteins from UniProt/SwissProt (119 sequences), UniProt/TrEMBL (2 sequences), and NCBI RefSeq (14 sequences), and for the SARS-CoV-2 and human proteins from UniProt/SwissProt (332 sequences) and from the supplementary material in [4] (26 sequences). We have taken the global alignment score between the amino acid sequences of two proteins, as implemented in BioPython [17], with a gap opening penalty of *−*7 and a gap extension penalty of *−*1, and normalized to [0, 1], as the similarity score between the proteins. In the protein sequences of P07203 and Q9BQE4, we substituted C (cysteine) for the rare amino acid U (selenocysteine). The corresponding integer linear programming problem instance has 83,628 variables, half of which are binary, and 84,069 constraints.

The alignment of the virus-host protein-protein interaction networks of SARS-CoV-1 and SARS-CoV-2 was computed with AMPL version 2018.10.22 [16] and Gurobi Optimizer version 8.1.0, using a personal computer with an Intel Core i7-8550U quad-core processor at 1.80 GHz and 32 GB of memory running Ubuntu 18.04 LTS. The optimal alignment was found in 517.35 seconds of AMPL time plus 3.16697 seconds of solver time for SARS-CoV-1 to SARS-CoV-2, and in 538.112 seconds of AMPL time plus 3.45882 seconds of solver time for SARS-CoV-2 to SARS-CoV-1. We set *λ* = 0.5 in both cases, and took the consensus between them as the alignment of the two virus-host protein-protein interaction networks.

We considered the host proteins that interact with the viral proteins in the consensus alignment, in each of the two networks. For these human proteins we obtained their Gene Ontology (GO) term annotations using GOnet [18], and measured the functional similarity between aligned human proteins with GOGO [19]. Tables 1 (structural proteins), 2–4 (non-structural proteins), and 5 (accessory proteins) show the alignment of viral proteins in the consensus alignment, along with the alignment of the human proteins they interact with, their molecular function ontology (MFO) score, their biological process ontology (BPO) score, and their cellular component ontology (CCO) score.

**Table 1.** Alignment of structural proteins in SARS-CoV-1 and SARS-CoV-2.

| SARS-CoV-1 | SARS-CoV-2 | BPO | CCO | MFO |
| --- | --- | --- | --- | --- |
| Spike protein S (P59594, P0DTC2) |  |  |  |  |
| O00303 | P48556 | 0.289 | 0.860 |  |
| Q9BYF1 | Q7L8L6 | 0.363 | 0.749 | 0.165 |
| O95295 | Q7Z5G4 | 0.485 | 0.855 |  |
| Envelope protein E (P59637, P0DTC4) |  |  |  |  |
| Q07817 | Q9Y3A6 | 0.299 | 0.806 |  |
| O00560 | Q6UX04 | 0.152 | 0.735 | 0.104 |
| Membrane protein M (P59596, P0DTC5) |  |  |  |  |
| O14920 | Q9UHD2 | 0.863 | 0.839 | 1.000 |
| P00403 | Q96HR9 | 0.077 | 0.816 |  |
| Q8TEB7 | Q96ER3 |  | 0.652 |  |
| O00303 | P48556 | 0.289 | 0.860 |  |
| P69849 | Q9NQC3 |  | 0.873 |  |
| Q9BYF1 | Q7L8L6 | 0.363 | 0.749 | 0.165 |
| Q03135 | Q00765 |  | 0.801 |  |
| Nucleocapsid protein N (P59595, P0DTC9) |  |  |  |  |
| P68104 | Q9HD40 | 0.625 | 0.748 | 0.511 |
| B8ZZN6 | Q9Y3U8 |  |  |  |
| Q92499 | Q9NR30 | 0.558 | 0.941 | 0.854 |
| Q53GL0 | Q8TAD8 | 0.131 | 0.851 |  |
| P09651 | Q8NCA5 | 0.090 | 0.692 | 0.631 |
| P14618 | Q13310 | 0.634 | 0.814 | 0.565 |
| P62937 | P67870 | 0.679 | 0.916 | 0.262 |
| B0QYN7 | P19784 |  |  |  |
| Q9Y4W2 | Q9NW13 | 0.622 | 0.917 | 0.674 |
| Q9HCD5 | P11940 | 0.055 | 0.869 | 0.868 |

**Table 2.** Alignment of non-structural proteins in SARS-CoV-1 and SARS-CoV-2.

| SARS-CoV-1 | SARS-CoV-2 | BPO | CCO | MFO |
| --- | --- | --- | --- | --- |
| Non-structural protein nsp1 (NP_828860, P0DTD1-PRO_0000449619) |  |  |  |  |
| O43447 | Q9H2H8 | 0.723 | 0.929 | 0.694 |
| P62937 | P67870 | 0.679 | 0.916 | 0.262 |
| P62942 | Q15370 | 0.470 | 0.863 | 0.083 |
| Q13427 | Q4V328 | 0.064 | 0.862 |  |
| Q9UKA8 | Q9Y680 | 0.049 | 0.848 | 0.209 |
| Q9Y4W2 | Q9NW13 | 0.622 | 0.917 | 0.674 |
| P28340 | P09884 | 0.887 | 0.965 | 0.792 |
| Non-structural protein nsp2 (NP_828861, P0DTD1-PRO_0000449620) |  |  |  |  |
| P05155 | Q9GZU3 |  |  |  |
| Q8WXF8 | Q99988 | 0.367 | 0.735 |  |
| Q7Z3Q1 | Q6NXT4 | 1.000 | 0.436 |  |
| Q7Z494 | Q08378 | 0.385 | 0.756 |  |
| P13796 | P52306 | 0.439 | 0.852 | 0.116 |
| P02768 | O14975 | 0.457 | 0.869 | 0.440 |
| Non-structural protein nsp4 (NP_904322, P0DTD1-PRO_0000449622) |  |  |  |  |
| P08949 | Q96DA6 | 0.372 | 0.539 |  |
| Q9Y4W2 | Q9NW13 | 0.622 | 0.917 | 0.674 |
| O95865 | Q9BSF4 | 0.092 | 0.772 |  |
| Non-structural protein nsp5 (NP_828863, P0DTD1-PRO_0000449623) |  |  |  |  |
| P23025 | Q8NEJ9 | 0.353 | 0.887 | 0.419 |
| P62942 | Q15370 | 0.470 | 0.863 | 0.083 |
| O75348 | O75347 | 0.104 | 0.713 | 0.201 |
| Q92802 | O95391 |  | 0.811 | 0.209 |
| Q9GZN8 | P82663 |  |  |  |
| Non-structural protein nsp6 (NP_828864, P0DTD1-PRO_0000449624) |  |  |  |  |
| O14498 | Q9BQQ3 | 0.407 | 0.792 |  |

**Table 3.** Alignment of non-structural proteins in SARS-CoV-1 and SARS-CoV-2.

| SARS-CoV-1 | SARS-CoV-2 | BPO | CCO | MFO |
| --- | --- | --- | --- | --- |
| Non-structural protein nsp7 (NP_828865, P0DTD1-PRO_0000449625) |  |  |  |  |
| A9UHW6 | Q8WVC6 |  |  | 0.141 |
| O95865 | Q9BSF4 | 0.092 | 0.772 |  |
| P13796 | P52306 | 0.439 | 0.852 | 0.116 |
| P49703 | P62330 | 0.464 | 0.749 | 0.852 |
| Q13564 | Q96K12 | 0.172 | 0.741 | 0.286 |
| Q53GL0 | Q8TAD8 | 0.131 | 0.851 |  |
| Q8TEB7 | Q96ER3 |  | 0.652 |  |
| Q92560 | Q96CN9 |  | 0.808 |  |
| P62258 | O00124 | 0.249 | 0.722 |  |
| Q9HCD5 | P11940 | 0.055 | 0.869 | 0.868 |
| A9UHW6 | Q8WVC6 |  |  | 0.141 |
| P08949 | Q96DA6 | 0.372 | 0.539 |  |
| P50583 | Q96A26 | 0.826 | 0.968 |  |
| P20290 | P51148 | 0.521 | 0.666 | 0.138 |
| P62879 | P62873 | 0.455 | 0.981 | 0.879 |
| Q99426 | P62820 | 0.289 | 0.761 |  |
| Q16548 | P61006 | 0.381 | 0.702 |  |
| Q92843 | P11233 | 0.485 | 0.727 |  |
| Q56VL3 | O43169 |  | 0.869 |  |
| Non-structural protein nsp8 (NP_828866, P0DTD1-PRO_0000449626) |  |  |  |  |
| P05155 | Q9GZU3 |  |  |  |
| P54274 | Q9H173 | 0.200 | 0.655 | 0.216 |
| P69849 | Q9NQC3 |  | 0.873 |  |
| P23588 | O00566 | 0.477 | 0.701 | 0.917 |
| Q9P0M6 | O14745 | 0.535 | 0.804 | 0.102 |
| P68104 | Q9HD40 | 0.625 | 0.748 | 0.511 |
| P49069 | Q13868 | 0.088 | 0.762 | 0.080 |
| Q96GS6 | Q9Y399 | 0.361 | 0.766 |  |
| P10415 | Q9NQT5 | 0.323 | 0.887 | 0.234 |
| P23025 | Q8NEJ9 | 0.353 | 0.887 | 0.419 |
| Q9Y2D1 | Q9H6F5 | 0.108 | 0.772 | 0.271 |
| Q9GZN8 | P82663 |  |  |  |
| Q13064 | O95260 | 0.714 | 0.234 | 0.083 |
| P23588 | O00566 | 0.477 | 0.701 | 0.917 |

**Table 4.** Alignment of non-structural proteins in SARS-CoV-1 and SARS-CoV-2.

| SARS-CoV-1 | SARS-CoV-2 | BPO | CCO | MFO |
| --- | --- | --- | --- | --- |
| Non-structural protein nsp9 (NP_828867, P0DTD1-PRO_0000449627) |  |  |  |  |
| P06733 | P31323 | 0.310 | 0.906 | 0.539 |
| Q5SQN1 | Q86VR2 | 0.112 | 0.371 |  |
| Q6P587 | P13984 | 0.255 | 0.828 | 0.457 |
| Q7Z494 | Q08378 | 0.385 | 0.756 |  |
| Q9BUV0 | P09601 |  |  |  |
| Q9UQN3 | O43633 | 0.932 | 0.885 |  |
| P25685 | Q9NZL9 | 0.179 | 0.912 | 0.879 |
| P09630 | Q15056 | 0.117 | 0.791 | 0.387 |
| Q9UMX0 | Q9BVL2 | 0.329 | 0.779 | 0.170 |
| Q6P587 | P13984 | 0.255 | 0.828 | 0.457 |
| Non-structural protein nsp10 (NP_828868, P0DTD1-PRO_0000449628) |  |  |  |  |
| P20290 | P51148 | 0.521 | 0.666 | 0.138 |
| P35354 | Q96AY3 | 0.100 | 0.895 | 0.607 |
| Q9Y2D1 | Q9H6F5 | 0.108 | 0.772 | 0.271 |
| Q8N488 | Q9HAV7 | 0.202 | 0.932 | 0.190 |
| Non-structural protein nsp12 (NP_828869, P0DTD1-PRO_0000449629) |  |  |  |  |
| P09630 | Q15056 | 0.117 | 0.791 | 0.387 |
| P30876 | Q66GS9 | 0.137 | 0.679 |  |
| P51948 | Q9H4P4 | 0.378 | 0.835 | 0.420 |
| Q5SQN1 | Q86VR2 | 0.112 | 0.371 |  |
| Q8TD31 | Q9UJC3 | 0.729 | 0.924 |  |
| O95299 | O14656 | 0.378 | 0.699 | 0.129 |
| Q99426 | P62820 | 0.289 | 0.761 |  |
| Q99471 | Q8N6S5 |  |  |  |
| Q9UQN3 | O43633 | 0.932 | 0.885 |  |
| O43447 | Q9H2H8 | 0.723 | 0.929 | 0.694 |
| P46379 | Q5T6F2 |  | 0.799 | 0.102 |
| Q9H000 | Q96IZ5 | 0.188 |  | 0.733 |
| Q92802 | O95391 |  | 0.811 | 0.209 |
| P98161 | O75592 | 0.379 | 0.725 | 0.596 |
| Non-structural protein nsp15 (NP_828872, P0DTD1-PRO_0000449632) |  |  |  |  |
| P62937 | P67870 | 0.679 | 0.916 | 0.262 |
| P51948 | Q9H4P4 | 0.378 | 0.835 | 0.420 |
| P49703 | P62330 | 0.464 | 0.749 | 0.852 |

**Table 5.** Alignment of accessory proteins in SARS-CoV-1 and SARS-CoV-2.

| SARS-CoV-1 | SARS-CoV-2 | BPO | CCO | MFO |
| --- | --- | --- | --- | --- |
| Accessory protein orf3a (P59632, P0DTC3) |  |  |  |  |
| Q13561 | Q8TD10 |  | 0.652 |  |
| P62258 | O00124 | 0.249 | 0.722 |  |
| Q03135 | Q00765 |  | 0.801 |  |
| Q99471 | Q8N6S5 |  |  |  |
| Q9BUV0 | P09601 |  |  |  |
| Accessory protein orf6 (P59634, P0DTC6) |  |  |  |  |
| P25685 | Q9NZL9 | 0.179 | 0.912 | 0.879 |
| Q13561 | Q8TD10 |  | 0.652 |  |
| Q5TBA9 | P49454 | 0.383 | 0.806 | 0.100 |
| P52292 | Q9NZJ7 | 0.152 | 0.808 |  |
| Q13287 | P78406 | 0.174 | 0.825 |  |
| Q13009 | P52948 | 0.169 | 0.829 | 0.170 |
| Accessory protein orf7a (P59635, P0DTC7) |  |  |  |  |
| P10415 | Q9NQT5 | 0.323 | 0.887 | 0.234 |
| P50583 | Q96A26 | 0.826 | 0.968 |  |
| Q07817 | Q9Y3A6 | 0.299 | 0.806 |  |
| Q16548 | P61006 | 0.381 | 0.702 |  |
| Q92843 | P11233 | 0.485 | 0.727 |  |
| O43765 | Q7Z4Q2 | 0.489 | 0.531 |  |
| Accessory protein orf9b (P59636, P0DTD2) |  |  |  |  |
| P08708 | Q9H773 | 0.509 | 0.861 | 0.185 |
| P25787 | Q9H2P9 | 0.319 | 0.701 | 0.120 |
| Q9UQN3 | O43633 | 0.932 | 0.885 |  |
| Q9P0M6 | O14745 | 0.535 | 0.804 | 0.102 |

We can observe that the four structural proteins in one network were aligned with the corresponding protein in the other network. Also, most of the non-structural proteins and half of the accessory proteins in one network where aligned with the corresponding protein in the other network. On the other hand, for each pair of aligned viral proteins, the highlighted proteins in the same column of a viral protein are the human proteins it interacts with. For instance, human proteins O00303 and Q9BYF1 interact with the SARS-CoV-1 spike protein P59594, while human protein Q7Z5G4 interacts with the SARS-CoV-2 spike protein P0DTC2. Table 1 shows that O00303 is aligned with P48556, Q9BYF1 is aligned with Q7L8L6, and O95295 is aligned with Q7Z5G4. Missing data is due to lack of GO term annotation for the two interacting proteins.

As can be seen in these tables, most of the aligned proteins have a cellular component ontology score above 0.700. This means that, despite the low number of conserved interactions, the aligned proteins share their cellular location. For instance, those human proteins that interact with the spike protein in SARS-CoV-1 are aligned with human proteins that interact with the membrane protein in SARS-CoV-2. However, some biological process ontology scores between aligned human proteins are very low. This can be explained by the lack of biological process ontology GO term annotation for one of the two interacting proteins.

With respect to molecular function ontology, it is remarkable that we obtained high scores for aligned proteins that interact with structurally different viral proteins. Indeed, one of the measures used to test the correctness of a protein-protein interaction network alignment is the *edge correctness* score, which measures the ratio of conserved edges in a given alignment. Edge correctness assumes that one of the aims of the alignment is to find similar regions between the two aligned networks, in terms of network topology. In the context of protein-protein interaction networks, it is also assumed that two proteins interact when they together carry out some biological function. For virus-host protein-protein interaction networks, viral proteins interact with host proteins to perturb the intracellular networks of their hosts to their advantage, and many virus-host interactions occur at the level of physical protein-protein interactions [7]. This means that a viral protein interacts with a host protein to carry out a cellular process, and this pathway of virus-host interactions constitutes the infection mechanism of the virus.

The question then arises, when a viral protein in one network is aligned with a viral protein in another network, should the host proteins that interact with one viral protein be aligned with those host proteins that interact with the other viral protein? Clearly the answer is yes, when the viral-host protein-protein interaction is a similar infectious process stage. Therefore, aligned virus-host interactions must entail conserved stages in the infectious process. However, non-conserved edges do not necessarily imply incorrect alignments. Indeed, when we analyze in more depth the functional description [3] in the UniProt database of the aligned human proteins that interact with Coronavirus proteins, we realise that they share a function related to viral infection, although their alignment introduces a non-preserved interaction. This is the case of the following pairs of proteins:

### O14920 and Q9UHD2

The molecular function ontology score of these proteins is 1.000. Human protein O14920 interacts with viral protein P59596 (membrane) of SARS-CoV-1, which is aligned with protein P0DTC5 (membrane) of SARS-CoV-2. On the other hand, human protein Q9UHD2 interacts with viral protein P0DTD1-PRO 0000449630 (orf9b) of SARS-CoV-2. However, the functional description of the aligned human proteins reflects correctness of the consensus alignment:

- O14920: Serine kinase that plays an essential role in the NF-kappa-B signaling pathway which is activated by multiple stimuli such as inflammatory cytokines, bacterial or viral products.
- Q9UHD2: Serine/threonine kinase that plays an essential role in regulating inflammatory responses to foreign agents. Following activation of toll-like receptors by viral or bacterial components, associates with TRAF3 and TANK and phosphorylates interferon regulatory factors (IRFs) IRF3 and IRF7 as well as DDX3X.

### Q92499 and Q9NR30

The molecular function ontology score of these proteins is 0.850. Human protein Q9NR30 interacts with viral protein P0DTC9 (nucleocapsid) of SARS-CoV-2, which is aligned with protein P59595 (nucleocapsid) of SARS-CoV-1. On the other hand, human protein Q92499 interacts with viral protein P0C6×7-PRO 0000037320 (proofreading exoribonuclease in replicase polyprotein 1ab) of SARS-CoV-1. The functional description of the aligned human proteins is:

- Q92499: Helicase required for Coronavirus IBV replication. Antiviral defense.
- Q9NR30: Component of a multi-helicase-TICAM1 complex that acts as a cytoplasmic sensor of viral double-stranded RNA (dsRNA) and plays a role in the activation of a cascade of antiviral responses including the induction of proinflammatory cytokines via the adapter molecule TICAM1.

### P49703 and P62330

The molecular function ontology score of these proteins is 0.850. Both are GTP-binding proteins. Human protein P49703 interacts with viral protein NP 828865 (nsp7) of SARS-CoV-1. Human protein P62330 interacts with viral protein P0DTD1-PRO0000449632 (nsp15) of SARS-CoV-2, which, as reported in [**?**], “has uridine-specific endoribonuclease (endoU) activity and is essential for viral RNA synthesis,” with the endoU domain being “one of the most conserved proteins among CoVs and related viruses, suggesting important functions in the viral replicative cycle.” The functional description of the aligned human proteins is:

- P49703: Small GTP-binding protein which cycles between an inactive GDP-bound and an active GTP-bound form, and the rate of cycling is regulated by guanine nucleotide exchange factors (GEF) and GTPase-activating proteins (GAP).
- P62330: GTP-binding protein involved in protein trafficking that regulates endocytic recycling and cytoskeleton remodeling. Activation is generally mediated by a guanine exchange factor (GEF), while inactivation through hydrolysis of bound GTP is catalyzed by a GTPase activating protein (GAP).

Therefore, it is not clear whether or not edge preservation should always be required in a correct alignment of virus-host protein-protein interaction networks. To reinforce this idea, we considered the functional similarity of all pairs of human proteins whose alignment would preserve edges, given the consensus alignment of 24 viral proteins. For each pair of aligned viral proteins (say, membrane proteins) we considered the biological process, the cell component, and the molecular function ontology scores of all pairs of human proteins that interact with the aligned viral proteins (say, all pairs of human proteins such that the first protein interacts with viral membrane protein P59596 of SARS-CoV-1 and the second protein interacts with viral membrane protein P0DTC5 of SARS-CoV-2). The cell component ontology score is above 0.800 for most of the aligned human proteins, but the highest molecular function ontology score is 0.852, while it is 1.000in the consensus alignment, and the highest biological process ontology score of the aligned human proteins is 0.670, while it is 0.863 in the consensus alignment.

Nevertheless, some of these pairs of human proteins whose alignment would preserve virus-host interactions do show high functional similarity scores, and it could be interesting to further study their role in the viral mechanism of host infection. Table 6 shows some of the highest ranking pairs of human proteins across biological process, cellular component, and molecular function ontology scores for the structural viral proteins in the consensus alignment, in descending order of average score. See S1 Appendix for more details.

**Table 6.** Human proteins that interact with SARS-CoV-1 and SARS-CoV-2 structural proteins, whose alignment would preserve virus-host interactions.

| SARS-CoV-1 | SARS-CoV-2 | BPO | CCO | MFO |
| --- | --- | --- | --- | --- |
| Spike protein S (P59594, P0DTC2) |  |  |  |  |
| Q9BYF1 | Q7Z5G4 | 0.520 | 0.859 | 0.137 |
| O00303 | Q7Z5G4 | 0.373 | 0.747 | 0.098 |
| O00303 | Q9C0B5 | 0.487 | 0.601 | 0.098 |
| Q9BYF1 | Q9C0B5 | 0.360 | 0.650 | 0.137 |
| Envelope protein E (P59637, P0DTC4) |  |  |  |  |
| Q07817 | O00203 | 0.548 | 0.805 | 1.000 |
| Q07817 | O60885 | 0.539 | 0.770 | 0.566 |
| O00560 | O00203 | 0.666 | 0.801 | 0.335 |
| Q07817 | P25440 | 0.402 | 0.759 | 0.397 |
| O00560 | Q8IWA5 | 0.531 | 0.864 | 0.105 |
| O00560 | P25440 | 0.353 | 0.734 | 0.335 |
| Membrane protein M (P59596, P0DTC5) |  |  |  |  |
| O14920 | Q96D53 | 0.310 | 0.889 | 0.852 |
| O14920 | Q9ULX6 | 0.246 | 0.880 | 0.638 |
| O14920 | P05026 | 0.472 | 0.743 | 0.549 |
| O14920 | P27105 | 0.473 | 0.712 | 0.574 |
| O14920 | Q7L8L6 | 0.308 | 0.865 | 0.449 |
| O14920 | P38606 | 0.284 | 0.842 | 0.491 |
| Nucleocapsid protein N (P59595, P0DTC9) |  |  |  |  |
| P09651 | Q8TAD8 | 0.630 | 0.891 | 0.906 |
| P09651 | P11940 | 0.436 | 0.954 | 0.906 |
| P09651 | Q6PKG0 | 0.572 | 0.786 | 0.906 |
| P14618 | P67870 | 0.693 | 0.909 | 0.657 |
| P09651 | Q9UN86 | 0.616 | 0.802 | 0.774 |
| P09651 | Q13310 | 0.425 | 0.855 | 0.906 |

We observed that, based on current knowledge, SARS-CoV-1 and SARS-CoV-2 share only 6 human proteins in their virus-host protein-protein interaction networks. On the one hand, aligned viral proteins in the consensus alignment obtained with our method show a sequence similarity of over 75% on the average, and most of the SARS-CoV-1 proteins are aligned with SARS-CoV-2 proteins that belong to the same category (spike, envelope, membrane, nucleocapsid, and the various non-structural and accessory proteins) in the genome organization of the viruses. On the other hand, the proposed alignment method does not preserve the virus-host interactions. This suggests that these viruses, despite their classification as human pathogens within the *Coronaviridae* family, do not follow the same detailed mechanism of host infection. We believe that further research on these aligned human proteins with high molecular function ontology scores, will help to elucidate the viral mechanism of infection and replication that is necessary to accomplish the goal of antiviral drug or vaccine discovery.

## Acknowledgments

We thank Francesc Rosselló for comments on an earlier version of this manuscript.

## Supporting information

### S1 Appendix. Human proteins that interact with SARS-CoV-1 and SARS-CoV-2 structural proteins, whose alignment would preserve

### virus-host interactions

For each pair of viral protein in the consensus alignment of the virus-host protein-protein interaction networks for SARS-CoV-1 and SARS-CoV-2, we show the molecular function ontology (MFO) score, the biological process ontology (BPO) score, and the cellular component ontology (CCO) score of the human proteins they interact with, in decreasing order of average score. Missing data is due to lack of GO term annotation for the two interacting proteins.

